# High-Yield Monocyte, Macrophage, and Dendritic Cell Differentiation from Induced Pluripotent Stem Cells

**DOI:** 10.1101/2021.04.29.441947

**Authors:** Lucas H. Armitage, Mohsen Khosravi-Maharlooei, Amy Meacham, Edward J. Butfiloski, Ryan Viola, Dieter Egli, Megan Sykes, Mark A. Wallet, Clayton E. Mathews

## Abstract

Differentiation of induced pluripotent stem cells (iPSC) into monocytes, monocyte-derived macrophages (MDM), and monocyte-derived dendritic cells (moDC) represents a powerful tool for studying human innate immunology and developing novel iPSC-derived immune therapies. Challenges include inefficiencies in iPSC-derived cell cultures, labor-intensive culture conditions, low purity of desired cell types, and feeder cell requirements. Here, a highly efficient method for differentiating monocytes, MDMs, and moDCs that overcomes these challenges is described. The process utilizes commercially-available materials to derive CD34^+^ progenitor cells that are apically released from a hemogenic endothelium. Subsequently, the hemogenic endothelium gives rise to highly pure (>95%), CD34^-^CD14^+^ monocytes in 19-23 days and yields 13.5-fold more monocytes by day 35 when compared to previous methods. These iPSC-monocytes are analogous to human blood-derived monocytes and readily differentiate into MDM and moDC. The efficient workflow and increase in monocyte output heightens feasibility for high throughput studies and enables clinical-scale iPSC-derived manufacturing processes.

## Introduction

Primary monocytes collected from anticoagulated human blood samples can be used directly in laboratory experiments or subjected to various differentiation conditions to create derivative populations of monocyte-derived macrophages (MDM) and monocyte-derived dendritic cells (moDC). Use of MDM and moDC is a straight-forward and widely-used approach to interrogating human innate antigen presenting cell (APC) and innate immune function. Blood is an easily accessed tissue that is readily available under informed consent (e.g. whole blood from research volunteers, buffy coats from blood banks, etc.). However, unlike T and B cells, monocytes do not expand after they are isolated from whole blood. In order to continue studying cells from the same donor continued blood draws are required; an effort that can prove difficult. This problem is compounded when studying rare genotypes where recruitment of sufficient donors is complex. Induced pluripotent stem cells (iPSC) and embryonic stem cells (ESC) offer a solution as differentiation of pluripotent stem cells to cells fo interest mitigates the need to recall donors. Further, rare genotypes can be introduced via genome engineering.

The introduction of ESC (Thomson et al., 1998) and later iPSC (Okita et al., 2007; Takahashi et al., 2007), was hailed as providing an unlimited source of human cells for research and therapeutics (Inoue et al., 2014). ESC and iPSC provide many advantages over primary cells; they can be indefinitely expanded (Takahashi et al., 2007), genetically manipulated (Ben Jehuda et al., 2018), and differentiated into rare cell types (e.g. CD34^+^ progenitor cells (Xu et al., 2012)) or cell types that occur in hard to access tissues (e.g. microglial cells (McQuade et al., 2018), pancreatic β cells (Pellegrini et al., 2018; Sui et al., 2018)). ESC and iPSC have great utility for disease modeling in isogenic systems, regenerative therapies, and drug discovery (Avior et al., 2016; Dimmeler et al., 2014). Recently, iPSC derived natural killer (NK) cells have been tested in phase 1 cancer immunotherapy clinical trials (NCT03841110, NCT04023071, and NCT04245722) paving the way for a potential new strategy to utilize renewable iPSC as a basis for groundbreaking therapies.

Protocols for differentiation of iPSC and ESC to monocytes, macrophages, and dendritic cells (DCs) have been previously described (Cao et al., 2019; Choi et al., 2009; van Wilgenburg et al., 2013; Yanagimachi et al., 2013). While these methods established the first successful protocols, there is room for improvement by increasing yield or eliminating the reliance on feeder cells that are essential for some protocols (Cao et al., 2019; Choi et al., 2009; van Wilgenburg et al., 2013; Yanagimachi et al., 2013). Here, we describe a modified method based on an earlier protocol (van Wilgenburg et al., 2013) that uses commercially available reagents to produce a meaningful improvement in yield and cell purity while maintaining the identity of the differentiated cells. We validate cell morphology, surface receptor expression, and function of monocytes, macrophages, and DC generated with the new protocol.

## Results

### iPSC exhibit normal karyotype and pluripotency markers

The iPSC lines, 2395 (Fredette et al., 2020), 1414 (Taylor et al., 2018), and 1-023 (Sui et al., 2018) were all previously subjected to quality control. Here, lines 1-039, 1-054, and 1-064, were assessed for normal karyotype and pluripotency. All three lines exhibited a normal karyotype when assessed with the G-banding method (Supplemental Figure 1A, 1B, and 1C respectively). In addition, lines 1-039, 1-054, and 1-064 expressed the pluripotency markers (OCT4, SSEA-4, and NANOG) when assayed via flow cytometry, (Supplemental Figure 1D, 1E, and 1F respectively).

### Rapid conversion of hematopoietic precursor producing hemogenic endothelium to high-yield CD14^+^ monocyte production

Our group has previously adopted an embryoid body-based differentiation protocol to generate iPSC-derived monocytes and macrophages (Taylor et al., 2018). These cells recapitulated critical aspects of normal human macrophage biology including supporting productive infection by HIV. However, the embryoid body (EB) step introduces a source of variability (many EBs fail to generate monocytes) and it presents a bottleneck for scalability. To overcome these challenges, we sought to derive a non-EB method for continuous monocyte differentiation from adherent progenitor cultures.

Here we utilized a commercially available differentiation kit that is intended for generation and collection of CD34^+^ hematopoietic stem cells (HSCs) for downstream applications (Figure 1A). The manufacturer’s protocol for the STEMDiff Hematopoietic Kit is geared towards collection of CD34^+^ hematopoietic precursors harvested on day 12 as the target population. Indeed, the cells that we collected on day 12 were uniformly CD34^+^ HSCs (Figure 1B). However, the adherent cell population that gives rise to the floating CD34^+^ HSCs is usually discarded once the HSCs have been collected. Instead, after removal of HSCs, the STEMdiff medium was replaced with monocyte factory medium (X-VIVO15 medium (Lonza, Morristown, NJ, USA), supplemented with 1X GlutaMAX^™^ (ThermoFisher Scientific, Waltham, MA, USA), 0.055 mM 2-mercaptoethanol (ThermoFisher), 100 ng/mL recombinant human M-CSF (PeproTech, Rocky Hill, NJ, USA) and 25 ng/mL recombinant human IL-3 (PeproTech)). Over the course of the next nine days (days 13-21 of the process), a change in the phenotype of collected floating cells occurred so that by day 21, more than 95% of harvested cells were CD14^+^CD34^-^ (Figure 1B). In this format, 65.7×10^6^ CD14^+^ monocytes were produced on average from a 24-well plate by day 35 while the orginal method only yielded 4.9×10^6^ monocytes from a single 6-well plate in our hands. The total average number of monocytes produced with this new method was 179×10^6^ by day 126 (Figure 1C; n = 3 (iPSC lines 2395, 1414, and 1-064)) compared to 12.4×10^6^ monocytes from the orginal method in our hands (Taylor et al., 2018; van Wilgenburg et al., 2013). This new method has two notable linear output phases after the initial 2-week burst of monocytes (65.7×10^6^ for days 21-35); the first averaged 12.2×10^6^ monocytes per week for days 35-70, and the second averaged 5.5×10^6^ monocytes per week for days 70-154 (Figure 1C). Upon examination of the hemogenic endothelium, it was observed that the CD14^+^ monocytes were released apically from large f-actin^+^ cells with morphology similar to that of cultured endothelial cells (Figure 1D).

**Figure 1.**
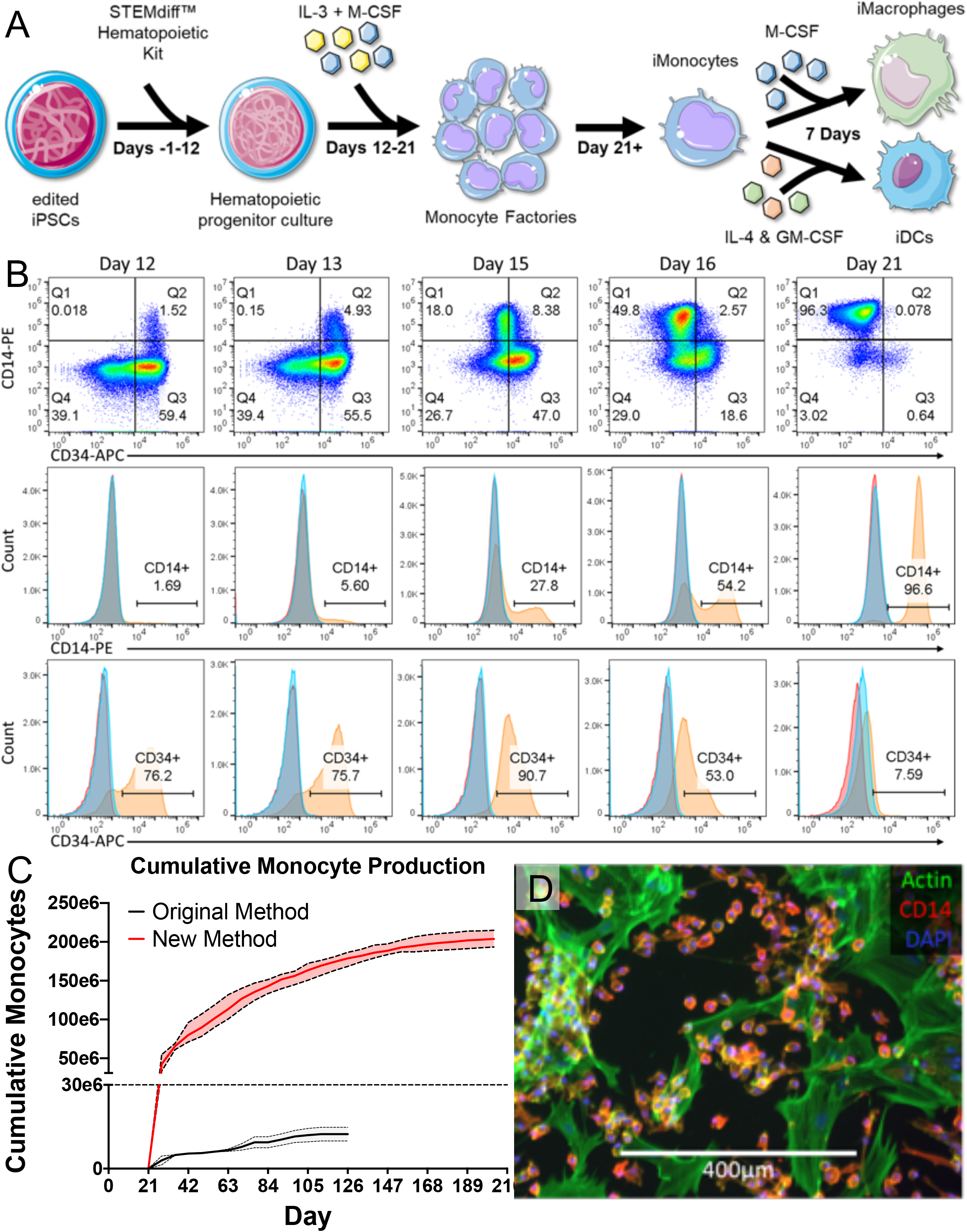
Differentiation of iPSC to monocytes, macrophages, and dendritic cells. A) Schematic of differentiation. iPSC are differentiated to a hematopoietic progenitor culture using the STEMdiff Hematopoietic Kit. The hematopoietic progenitors are harvested and the remaining hemogenic endothelium is cultured in the presence of rhIL-3 and rhM-CSF and starts producing monocytes. The monocytes can be cultured in the presence of rhM-CSF to differentiate into MDM or rhIL-4 and rhGM-CSF to differentiate into moDC. B) Cells produced during the conversion of the hemogenic endothelium from CD34^+^ hematopoietic cell production to CD14^+^ monocyte production stained with αCD14-PE and αCD34-APC and analyzed via flow cytometry. C) Cumulative monocyte production of this method (red, n=6) vs. a previously published method (van Wilgenburg et al., 2013) (black, n=2). D) A monocyte producing hemogenic endothelium was stained for Actin (Green), CD14 (Red), and nuclei (Blue, DAPI) and imaged on an EVOS FL Imaging System.

### iPSC and ESC-derived and primary monocytes exhibit similar morphology and marker expression

To learn if iPSC (lines 1-039, 2395, and 1-064) and ESC-derived (line MEL-1) monocytes were similar to their primary counterparts, cell morphology was assessed via fluorescence microscopy and cell surface marker expression was assessed via flow cytometry. iPSC and ESC-derived monocytes were similar to classical blood monocytes in morphology and surface marker expression (Ziegler-Heitbrock et al., 2010). The monocytes were round with large nuclei and expressed similar amounts of CD14, CD64, and CD16 when compared to primary monocytes (Figure 2A, B, and C). The original method resulted in noticeably larger iPSC-monocytes when compared to primary monocytes (Taylor et al., 2018). The method described here also yields iPSC-derived monocytes that are significantly larger than primary monocytes (Figure 2D; iPSC-derived monocytes averaged 10.06 μm and primary monocytes averaged 7.68 μm in diameter, **p = 0.0014 in unpaired Student’s t-test). One important note is that these iPSC-derived cells exhibited higher autofluorescence than their primary cell derived counterparts. This autofluorescence is readily evident in the representative histograms when comparing unstained cells (Figures 2B, 3B, and 4B).

**Figure 2.**
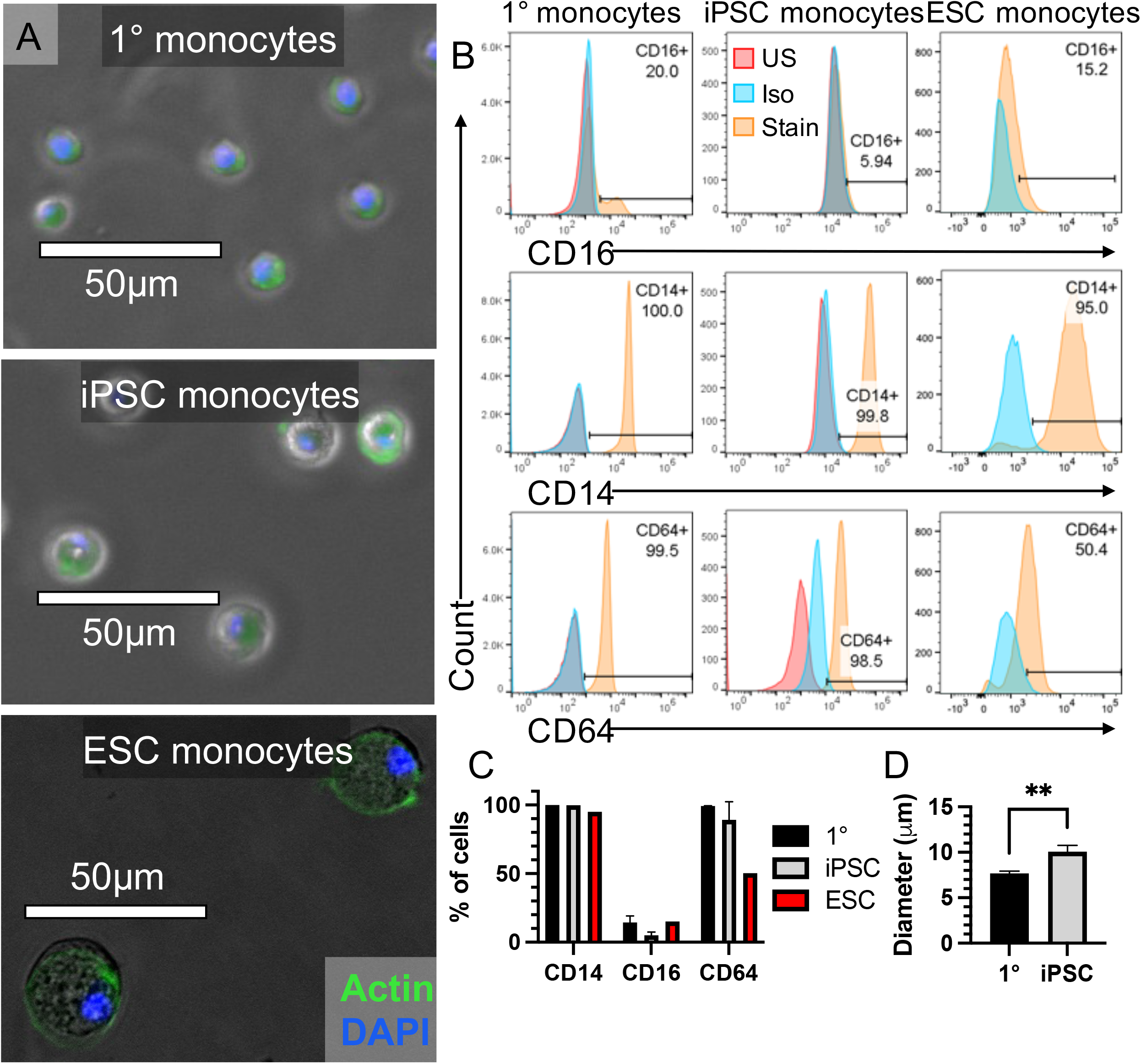
iPSC and ESC-derived monocytes are similar to primary classical monocytes in both morphology and surface marker expression. A) Primary and iPSC-derived monocytes were stained with actin green and DAPI and imaged on an EVOS FL Imaging System. ESC-derived monocytes were also stained with actin green and DAPI and imaged on a Leica Fluorescent DMI 6000B wide field microscope. B) Representative histograms of surface marker expression on primary, iPSC, and ESC-derived monocytes stained with αCD16-FITC, αCD14-PE, and αCD64-APC then analyzed via flow cytometry. C) Surface marker expression quantified by percent of monocytes expressing CD14, CD16, and CD64. Primary monocyte and iPSC-derived monocytes, n=3; ESC-derived monocytes, n=1. D) Diameter of primary and iPSC-derived monocytes.

**Figure 3.**
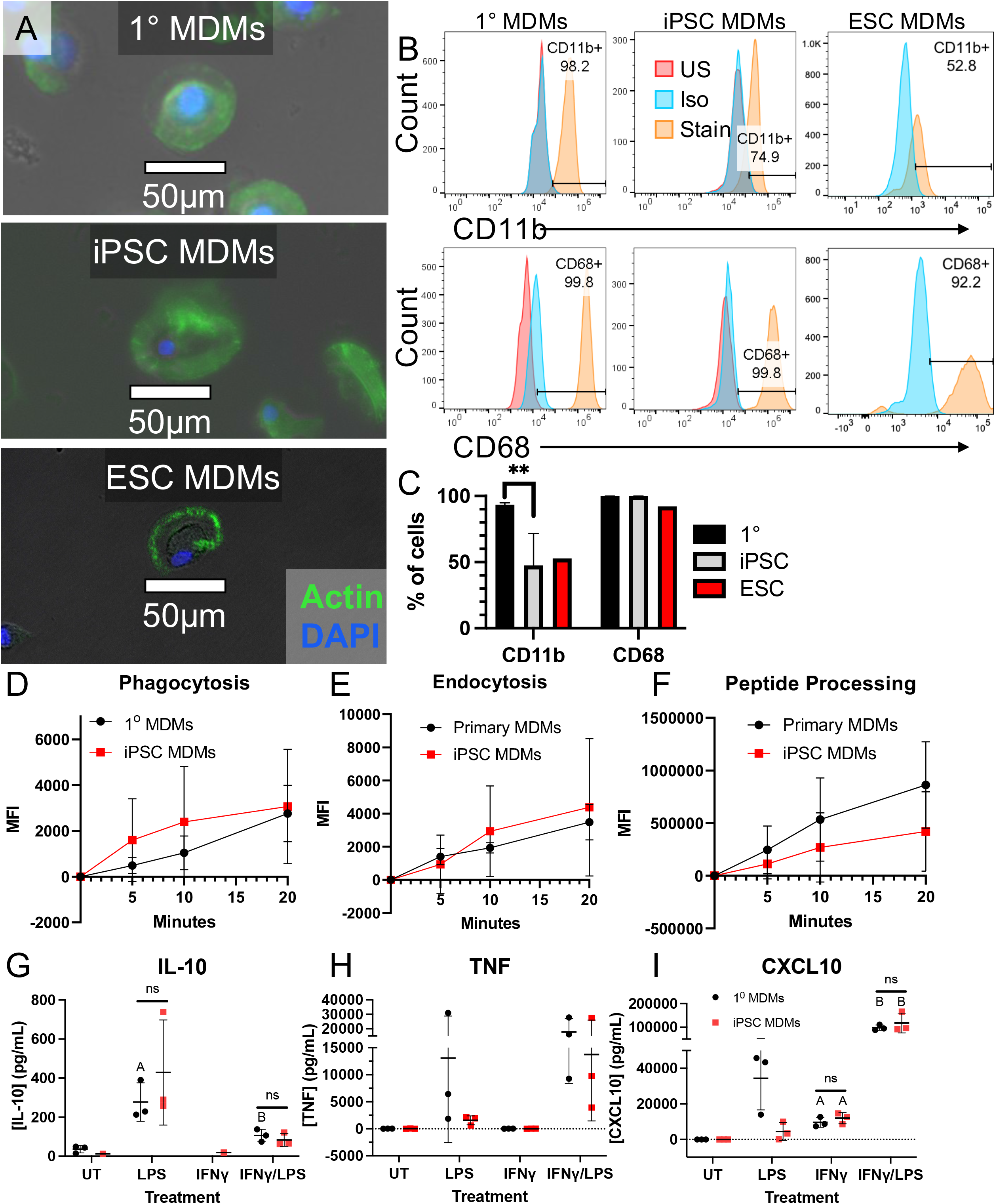
iPSC and ESC-derived MDM are similar to primary MDM in morphology, surface marker expression, and function. A) Primary and iPSC-derived MDM were stained with actin green and DAPI and imaged on an EVOS FL Imaging System. ESC-derived MDM were also stained with actin green and DAPI and imaged on a Leica Fluorescent DMI 6000B wide field microscope. B) Representative histograms of surface marker expression on primary, iPSC, and ESC-derived MDM stained with αCD11b-FITC and αCD68-PE and analyzed via flow cytometry. C) Surface marker expression quantified by percent of MDM expressing CD11b, and CD68. Primary MDM and iPSC-derived MDM, n=3; ESC-derived MDM, n=1. D-F) Uptake of D) pHrodo Red *S. aureus* bioparticles, E) pHrodo Red labeled dextran, and F) uptake and proteolysis of DQ Ovalbumin measured via flow cytometry. The data were analyzed via 2-way ANOVA and no significant differences were detected between iPSC-derived and primary MDM. F) ELISA for IL-10. IL-10 was not detected in some of the untreated and the IFNγ-treated groups. The data were analyzed via multiple T tests and there were no significant differences between iPSC-derived and primary MDM. Letters on the graph indicate significant difference compared to untreated of the same group. G) ELISA for TNF. The data were analyzed via 2-way ANOVA and no significant differences were detected between iPSC-derived and primary MDM. Letters on the graph indicate significant difference compared to untreated of the same group.H) ELISA for CXCL10. The data were analyzed via 2-way ANOVA and no significant differences were detected between iPSC-derived and primary MDM. Letters on the graph indicate significant difference compared to untreated of the same group.

**Figure 4.**
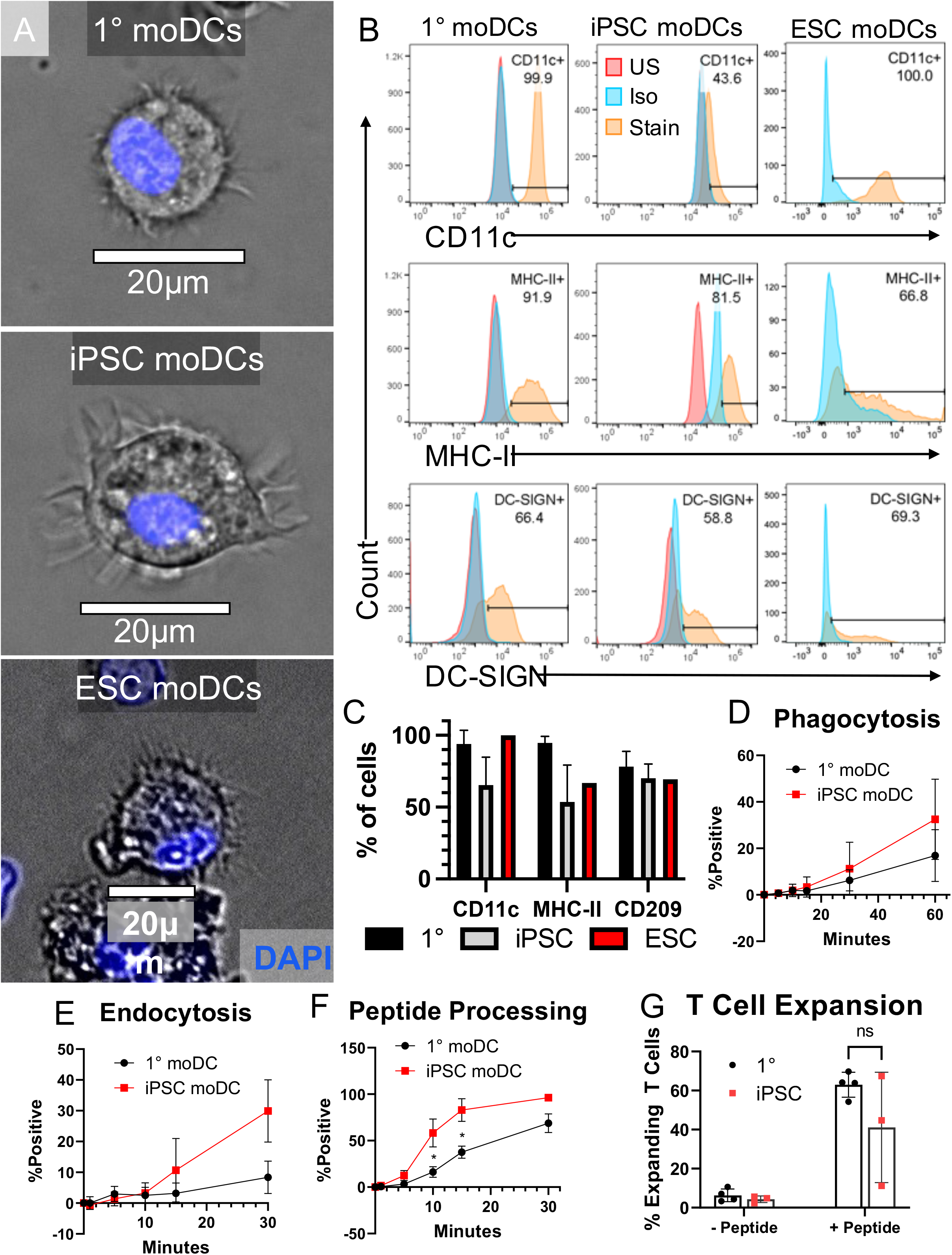
iPSC and ESC-derived moDC are similar to primary moDC in morphology, surface marker expression, and function. A) Primary and iPSC-derived moDC were stained with DAPI and imaged on an EVOS FL Imaging System. ESC-derived moDC were also stained with DAPI and imaged on a Leica Fluorescent DMI 6000B wide field microscope. B) Representative histograms of surface marker expression on iPSC and primary moDC stained with αCD11c-FITC, αHLA-DR, DP, DQ-PE/Cy7 and analyzed via flow cytometry. C) Surface marker expression quantified by percent of moDC expressing CD11c, MHC-II, and CD209. Primary moDC and iPSC-derived moDC, n=3; ESC-derived moDC, n=1. D-E) Uptake of D) pHrodo Red *S. aureus* bioparticles, E) pHrodo Red labeled dextran, and D) uptake and proteolysis of DQ Ovalbumin measured via flow cytometry. G) Quantification of expansion of MART1-CD8^+^ T cell avatars under different moDC loading conditions.

### iPSC-, ESC-derived, and primary MDM exhibit similar morphology, marker expression, and function

To determine if iPSC (lines 1414, 2395, and 1-064) and ESC-derived (line MEL-1) MDM were similar to primary MDM, cell morphology was assessed via fluorescence microscopy and cell surface marker expression was assessed via flow cytometry. iPSC and ESC-derived MDM were similar to primary MDM in morphology and surface marker expression; they were adherent with the majority of cells having a large cytoplasm and a centered nucleus, considered the classical “fried egg” morphology (Figure 3A) (Young et al., 1990). iPSC and ESC-derived MDM expressed similar levels of CD68 to primary MDM however iPSC-MDM expressed significantly lower levels of CD11b (Figure 3B and C, **p = 0.0097 in multiple comparisons of 2-way ANOVA).

A hallmark of macrophages is their ability to phagocytose and digest invading pathogens as well as their ability to endocytose and process antigen for presentation on major histocompatibility molecules to T cells. To verify that iPSC-derived MDM had a similar phagocytic capacity compared to primary MDM, both cell types were incubated with pHrodo Red *S. aureus* bioparticles. pHrodo Red dye is weakly fluorescent at neutral pH and this fluorescence increases as pH decreases, making it ideal for tracking acidification of lysosomes. In the current system, as the pHrodo Red *S. aureus* bioparticles are phagocytosed the formation of the phagolysosome or lysosome occurs and these vesicles undergo rapid acidification leading to increased fluorescence (Gordon, 2016). iPSC-derived and primary MDM phagocytosed and acidified phagosomes at similar rates as indicated by the increase in fluorescence of cells incubated with pHrodo Red *S. aureus* bioparticles (Figure 3D). To confirm that iPSC-derived MDM are comparable to primary MDM in their ability to endocytose, both cell types were loaded with pHrodo Red dextran, which is taken up via endocytosis. iPSC-derived and primary MDM endocytosed and acidified endocytic vesicles at similar rates as indicated by the increased fluorescence of cells incubated with pHrodo Red dextran (Figure 3E). The ability to uptake and process antigen was assessed by incubating iPSC-derived and primary MDM with DQ Ovalbumin (DQ-OVA), a self-quenched conjugate of ovalbumin that becomes fluorescent when it undergoes proteolysis. This is process is the same for uptaking and loading foreing antigen onto MHC molecules (Ciudad et al., 2017; Collin and Bigley, 2018). iPSC-derived MDM processed DQ-OVA at a similar rate to primary MDM (Figure 3F).

As one of the front-line sentinels of the immune system, macrophages are able to eliminate bacteria while simultaneously limiting tissue damage; one well-known way this balancing act is accomplished is through secretion of TNF and IL-10 in response to TLR agonists (e.g. LPS, CpG DNA, diacyl and triacyl lipopeptides, etc.) (Couper et al., 2008; Kang et al., 2019). TNF supports innate immune system activation and control of invading bacteria while IL-10 helps limit tissue damage (Kang et al., 2019). When macrophages are unable to control bacterial infections on their own, as is the case for *mycobacterium tuberculosis*, IFNγ from T cells helps fully activate macrophages to destroy invading pathogens (Couper et al., 2008). IFNγ inhibits IL-10 secretion from macrophages while augmenting TNF and CXCL10 production (Kang et al., 2019). These responses are well characterized and represent another hallmark function of macrophages. Upon LPS stimulation, both iPSC-derived and primary MDM upregulated IL-10 and TNF but not CXCL10 secretion (Figure 3G, H, and I). Treatment with recombinant human IFNγ (rhIFNγ) alone did not induce IL-10 or TNF expression but it did induce CXCL10 secretion (Figure 3G, H, and I). Upon treatment with rhIFNγ and LPS, both iPSC-derived and primary MDM downregulated IL-10 secretion and further upregulated TNF and CXCL10 secretion when compared to LPS treatment alone (Figure 3G, H, and I). There were no significant differences when comparing the response of iPSC-derived MDM to primary MDM in these assays. Thus iPSC-derived MDM and primary MDM are similar in morphology, cell marker expression, and function.

### iPSC-, ESC-derived, and primary moDC are similar in morphology, surface marker expression, and function

To verify if iPSC-derived (lines 1414, 2395, and 1-064) and ESC-derived (line MEL-1) moDC were similar to primary moDC, cell morphology was assessed via fluorescence microscopy and cell surface marker expression was assessed via flow cytometry. iPSC and ESC-derived moDC were similar to primary moDC in morphology and surface marker expression (Figure 4A and B). iPSC-, ESC-derived and primary moDC were lightly adherent, round, and had small dendrites (Figure 4A). The iPSC and ESC-derived moDC expressed CD11c (Verreck et al., 2006), MHC-II (Collin and Bigley, 2018; Sim et al., 2016), and CD209 (DC-SIGN) (Collin and Bigley, 2018; Sim et al., 2016) at levels similar to primary moDC (Figure 4B and C).

One of the well-defined features of moDC is their ability to uptake and process antigen for presentation to T cells and to subsequently induce expansion of an effector T cell population (Liu et al., 2019). iPSC-derived and primary moDC phagocytosed pHrodo Red *S. aureus* bioparticles (Figure 4D) and endocytosed pHrodo Red Dextran at similar rates (Figure 4E). iPSC-moDC outperformed primary-moDC at peptide processing at 10 and 15 minutes, however, by experimental endpoint (30 minutes) there was no significant difference between the two groups (Figure 4F). When iPSC-derived and primary moDC were left unloaded and co-cultured with MART1-CD8^+^ T cell avatars they induced little T cell expansion (Figure 4G). When iPSC-derived and primary moDC were loaded the MART-1 peptide recognized by the MART1 TCR and co-cultured with MART1-CD8^+^ T cell avatars they induced similar amounts of T cell expansion in an antigen specific manner (Figure 4G). Thus these iPSC and ESC-derived moDCs are a good model for primary moDCs.

## Discussion

Currently, it is possible to differentiate iPSC and ESC to monocytes, however, existing protocols result in low yields, involve labor intensive cell culture steps, rely on feeder cells, and require purification steps (Cao et al., 2019; Choi et al., 2009; van Wilgenburg et al., 2013; Yanagimachi et al., 2013). Indeed the limiting steps in the original protocol our group utilized were waiting for EBs to attach and start producing monocytes, which did not always occur (Taylor et al., 2018). Here we present a new high-yield iPSC to monocyte differentiation protocol that produces ~65 x 10^6^ monocytes in 35 days and has two linear output phases; days 35-70 – 12.2×10^6^ per week and days 70-142 – 5.5×10^6^ per week (Figure 1C). This is more than a 3-fold increase by day 35 compared to previously published methods (Cao et al., 2019; Choi et al., 2009; van Wilgenburg et al., 2013; Yanagimachi et al., 2013). Further, the procedure does not require additional sorting or purification as there are no feeder cells and the harvested cells are >95% CD14^+^ monocytes. The monocytes, MDM, and moDC are comparable in form and function to those attained with other protocols. The increase in monocyte yields and linear monocyte production phases with this protocol adds to the utility of iPSC-derived monocytes, macrophages, and dendritic cells.

Many experiments that utilize monocytes, MDM, or moDC require high numbers of cells. For example, adoptive transfer of monocytes into mice can require 30e6 monocytes per mouse and equate to over a half a billion cells in small experiments with 2 or more study groups of reasonable size (Cao et al., 2019; Choi et al., 2009; van Wilgenburg et al., 2013; Yanagimachi et al., 2013). Such experiments would require current monocyte differentiation protocols to be scaled up by at least 16-fold, however the protocol described here can fulfill these requirements with only three 12-well plates. Adoptive transfer experiments are not the only experiments that will benefit from increased yields; any experiments requiring high numbers of monocytes, MDM, or moDC (e.g. drug toxicity screenings for monocytes, MDM, and moDC, liver drug toxicity screenings where monocytes and MDMs play a role in propagating acute liver damage, etc.) would benefit from the implementation of this protocol.

Not only does this method produce far more monocytes than other methods but it is also more easily implemented. A commercially available kit from Stem Cell Technologies, the STEMdiff Hematopoietic Kit, has been used to standardize the initial steps (Figure 1A). This kit produces CD34^+^CD45^+^ hematopoietic progenitors in a 2D culture, eliminating the need to form embryoid bodies (EBs) and the labor-intensive culture of the EBs while they attach to culture ware. The increased ease in differentiating iPSC to monocytes will elevate reproducibility. An additional benefit to using this kit is the byproduct produced, iPSC-derived CD34^+^ hematopoietic stem cells, which can be differentiated into other cell types for studies such as NK cells (Woll et al., 2009), embryonic red blood cells (Olivier et al., 2006), microglial cells (McQuade et al., 2018), B cells (Carpenter et al., 2011). This new method is also highly reliable, producing many monocytes from multiple iPSC donors (n=6) and an ESC line reprogrammed using different techniques (Table 1). Like classical blood monocytes, these monocytes are CD14^+^, CD64^+^, and CD16^-^ (Figure 1B and Figure 2). The monocytes readily differentiate into functional macrophages (Figure 3) and dendritic cells (Figure 4) that are similar in surface marker expression, morphology, and function to their primary cell derived counterparts. Importantly, the monocytes produced here are similar to previous iPSC-derived monocytes in quality. These monocytes, MDM, and moDC are very similar to their primary counterparts; however, there are some notable differences.

**Table 1.** iPSC and ESC lines used

| Line | Reprogramming method | Cell source | Institution |
| --- | --- | --- | --- |
| 2395 | Non-integrating sendai vector | CD34<sup>+</sup> peripheral stem cells | UF |
| 1414 | Non-integrating sendai vector | CD34<sup>+</sup> peripheral stem cells | UF |
| 1-023 | Retroviral vector | Fibroblasts | Columbia |
| 1-039 | Transient mRNA transfection | Fibroblasts | Columbia |
| 1-054 | Transient mRNA transfection | Fibroblasts | Columbia |
| 1-064 | Transient mRNA transfection | Fibroblasts | Columbia |
| MEL-1 | N/A | ESC | N/A |

The iPSC-derived monocytes are larger (Figure 2A and D) when compared to primary monocytes. One possible cause of the size difference is the method of culture. Monocytes arise in vivo from the bone marrow and enter circulation where they are subjected to mechanical stresses (i.e. shear stress, pressure). However, the culture method described here is a static culture method where the monocytes never enter circulation and are not subject to arterial/venous pressures or shear stress. Shear stress, can directly influence cell size (Heo et al., 2012; Holtzclaw et al., 2010) and might explain this observed difference. Another difference observed is the decreased expression of CD11b on iPSC-derived MDM compared to primary MDM (Figure 3B). The iPSC-derived monocytes, MDM, and moDC exhibit increased autofluorescence compared to their primary cell counterparts possibly due to their increased size (Figure 3B: MFI Unstained primary MDM 19287 ± 1759 vs. iPSC MDM 59604 ± 3814 when excited with the 488nm laser and detected with a 533/30 band-pass filter). This is most prominent when analyzing via flow cytometry and utilizing fluorophores that are excited with the 488nm laser and detected 500-625nm range, such as FITC or PE. The antibody used to detect CD11b (Figure 3B) was conjugated to FITC (Table 2). The increased autofluorescence of the iPSC-derived cells masks differences between the stained and unstained thus decreasing the detection of CD11b (Figure 3B).

**Table 2.** Antibodies and isotype controls for flow cytometry.

| Antigen | Fluorophore | Source | Clone | Isotype | Company | Catalog # |
| --- | --- | --- | --- | --- | --- | --- |
| OCT4 | AF488 | mouse | 3A2A20 | IgG2b, κ | BioLegend | 653706 |
| SSEA-4 | PE | mouse | MC-813-70 | IgG3, κ | BioLegend | 330406 |
| NANOG | AF647 | mouse | 16H3A48 | IgG1, κ | BioLegend | 674210 |
| CD14 | PE | mouse | M5E2 | IgG2a, κ | BioLegend | 301806 |
| CD34 | APC | mouse | 581 | IgG1, κ | BioLegend | 343510 |
| CD16 | FITC | mouse | eBioCB16 | IgG1, κ | ThermoFisher | 11-0168-42 |
| CD64 | APC | mouse | 10.1 | IgG1, κ | ThermoFisher | 17-0649-42 |
| CD11b | FITC | mouse | ICRF44 | IgG1, κ | BioLegend | 301330 |
| CD68 | PE | mouse | eBioY1/82A | IgG2b, κ | ThermoFisher | 12-0689-41 |
| CD11c | APC | mouse | Bu15 | IgG1, κ | BioLegend | 337208 |
| CD11c | FITC | Mouse | Bu15 | IgG1, κ | BioLegend | 337214 |
| HLA-DR, DP, DQ | APC | mouse | Tü39 | IgG2a, κ | BioLegend | 361714 |
| HLA-DR, DP, DQ | PE-Cy7 | mouse | Tü39 | IgG2a, κ | BioLegend | 361708 |
| CD209 | APC | rat | eB-h209 | IgG2a, κ | ThermoFisher | 17-2099-42 |
| Isotype Control | AF488 | mouse | MPC-11 | IgG2a, κ | BioLegend | 400329 |
| Isotype Control | PE | mouse | MG3-35 | IgG3, κ | BioLegend | 401320 |
| Isotype Control | AF647 | mouse | MOPC-21 | IgG1, κ | BioLegend | 400136 |
| Isotype Control | PE | mouse | MOPC-173 | IgG2a, κ | BioLegend | 400214 |
| Isotype Control | APC | mouse | MOPC-21 | IgG1, κ | BioLegend | 400122 |
| Isotype Control | FITC | mouse | MOPC-21 | IgG1, κ | BioLegend | 400109 |
| Isotype Control | PE | mouse | eBMG2b | IgG2b, κ | ThermoFisher | 12-4732-42 |
| Isotype Control | PE/Cy7 | mouse | MOPC-173 | IgG2a, κ | BioLegend | 400232 |
| Isotype Control | APC | rat | eBR2a | IgG2a, κ | ThermoFisher | 17-4321-81 |

An additional difference between the iPSC-derived cells and primary cells was in moDC, where the uptake and processing of DQ-ova was more rapid in iPSC-derived moDC compared to primary moDC (Figure 4F). While the iPSC-derived moDC processed DQ-ova faster than primary moDC (Figure 4F; 10 minute and 15 minute time points), by experimental endpoint (Figure 4F; 30 minute time point) the difference was no longer significant. One possible reason the iPSC-derived moDC uptake and process peptide faster is that they are larger and have more surface area to internalize DQ-ova through (Figure 4A). Notably, while there was not a significant difference in the ability of iPSC-derived moDC to expand CD8+ T cell avatars compared to primary moDC, one line of iPSC-derived moDC barely induced any expansion when peptide was present (Figure 4G). While two of the iPSC lines utilized here (1-039 and 2395) expanded CD8^+^ T cells similarly to the primary moDC, the third line, 1-023, was a poor stimulator of T cell proliferation. Upon karyotyping we found line 1-023 has an abnormal karyotype, 46,XY,t(1;12)(p36.2;q24.1)[20] (Supplemental Figure 2A; indicated by the blue arrows). This abnormality arose in culture as the original 1-023 iPSC line karyotype was normal (Supplemental Figure 2B). It is possible that this chromosomal abnormality has impaired the ability of moDC derived from iPSC line 1-023 to present antigen. While there are minor differences observed between these iPSC-derived cells and their primary counterparts, overall, the cells share morphology, marker expression, and, most importantly, function.

In conclusions, we have developed a new high-yield method for differentiating iPSC to monocytes, MDM, and moDC adding a powerful tool to the growing protocols for differentiating iPSC to relevant cells/tissues for the study of human health and disease.

## Acknowledgements

The University of Florida Center for Immunology and Transplantation, The University of Florida Center for Cellular Reprogramming, The University of Florida Diabetes Institute, and The University of Florida Clinical and Translational Science Institute, and the Columbia Center for Translational Immunology (CCTI) Flow Cytometry Core facility (supported by the Office of the NIH Director (S10OD020056, S10RR027050, P30CA013696, 5P30DK063608, and R01DK106436)) were all essential to the completion of these studies.

This work was supported by research grants from the National Institutes of Health UC4 DK104194 (CEM), R01 DK127497 (CEM), UG3 DK122638 (CEM), P01 AI042288 (CEM), T32 DK108736 (LHA), R24 GM119977, T32 DK108736 (LHA), NIAID P01 AI04589716 (MS), UC4 DK104207 (MS), Leona and Harry Helmsley Charitable Trust for the collection of T1D and control cell lines (D.E.), and the Sebastian Family Endowment for Diabetes Research (CEM).

## Author Contributions

LHA, CEM, and MAW conceived of the idea, designed the research plan, performed experiments, analyzed data, interpreted results of experiments, and participated in all aspects of writing the manuscript. MK-M performed experiments, analyzed data, interpreted results of experiments, and edited the manuscript. EB performed experiments, analyzed data, interpreted results of experiments, and edited the manuscript. AM, RV and DE generated iPSC, contributed to discussion, and edited the manuscript. MS contributed to research design as well as discussion, and edited the manuscript. All authors contributed to the article and approved the submitted version.

---

Supplemental Figure 1. Quality control for iPSC lines 1-039, 1-054, and 1-064. A) Line 1-039 displays a normal karyotype, 46XX. B) Line 1-059 displays a normal karyotype, 46XX. C) Line 1-064 displays a normal karyotype, 46XX. D) Line 1-039 expresses the pluripotency markers OCT4, SSEA-4, and NANOG. E) Line 1-054 expresses the pluripotency markers OCT4, SSEA-4, and NANOG. F) Line 1-064 expresses the pluripotency markers OCT4, SSEA-4, and NANOG.

Supplemental Figure 2. Karyotypes from line 1-023. A) The abnormal karyotype of the current line. B) The normal karyotype of the original line.

## Materials and Methods

### iPSC and ESC Maintenance and Quality Control

All critical reagents for this protocol are listed in Table 3. iPSC were maintained as previously described (Cao et al., 2019; Choi et al., 2009; van Wilgenburg et al., 2013; Yanagimachi et al., 2013). Briefly, iPSC were cultured on Matrigel-(Corning, Corning, NY, USA) coated 6-well plates in mTeSR Plus (STEMCELL Technologies, Vancouver, BC, Canada) with medium changes every other day. ESC were maintained on Matrigel coated plates in mTeSR1 (STEMCELL Technologies) with daily medium changes. Both cell types were passaged every 7 days or as they reached 70-80% confluency using ReLeSR (STEMCELL Technologies). All iPSC and ESC lines used for these studies are listed in Table 1 along with the method of reprogramming and institution they were reprogrammed at. The ESC line, MEL-1 (NIH Approval # NIHhESC-11-0139) and iPSC lines from donors 2395, 1414, 1-023, 1-039, 1-054, and 1-064 were used. Creation of the iPSC lines from donors 2395 and 1414 was previously described (Fredette et al., 2020; Taylor et al., 2018). iPSC lines 1-023, 1-039, 1-054, and 1-064 were kindly provided by the New York Stem Cell Foundation and Columbia University. Line 1-023 was reprogrammed from fibroblast via retroviral transfection and was previously described (Sui et al., 2018; Wang et al., 2015). Lines 1-039, 1-054, and 1-064 were reprogrammed from fibroblast via transient mRNA transfection as previously described (Johannesson et al., 2014; Sui et al., 2018; Yamada et al., 2014). The karyotyping for lines 1-039 and 1-054 were performed by UF Health Pathology Laboratories (Gainesville, FL) via the G-banding method. The karyotype for line 1-064 was performed by Cell Line Genetics (Madison, WI, USA) also using the G-banding method. Lines 1-039, 1-054, and 1-064 were also analyzed via flow cytometry for expression of the pluripotency markers, OCT4, SSEA-4, and NANOG (Table 2). iPSC were dissociated to single cells using ACCUTASE (StemCell Technologies) and pelleted at 300xg for 5 minutes at room temperature. iPSC were then fixed/permeabilized and stained using the Fixation/Permeabilization Solution Kit (BD Biosciences, Franklin Lakes, NJ, USA) according to the manufacturer’s protocol. Stained iPSC were then analyzed on an Accuri C6 (BD Biosciences) and data were analyzed in FlowJo Version 10 (BD Biosciences).

**Table 3.** Critical reagents for iPSC to monocyte, MDM, and moDC differentiation.

| Item | Company | Catalog # |
| --- | --- | --- |
| CryoStor® CS10 | Stemcell Technologies | 7930 |
| Corning® Matrigel® hESC-Qualified Matrix,<br>LDEV-free, 5 mL | Corning | 354277 |
| DMEM/Hams F-12, 50/50, 1X | Corning | 10-103-CV |
| mTeSR™ Plus cGMP, stabilized feeder-free<br>maintenance medium for human ES and iPS<br>Cells | Stemcell Technologies | 05825 |
| ReLeSR™ | Stemcell Technologies | 05872 |
| 70µM nylon cell strainer | Fisher Scientific | 22-363-548 |
| STEMdiff™ Hematopoietic Kit | Stemcell Technologies | 05310 |
| X-VIVO™ 15 Serum-free Hematopoietic Cell<br>Medium | Lonza | 04-418Q |
| 2-Mercaptoethanol (55mM) | Fisher Scientific | 21-985-023 |
| GlutaMAX™ Supplement | Fisher Scientific | 35-050-061 |
| recombinant human M-CSF | PeproTech | 300-25 |
| recombinant human IL-3 | PeproTech | 200-03 |
| Costar® 6-well Clear Flat Bottom Ultra-Low<br>Attachment Multiple Well Plates | Corning | 3471 |
| DMEM w/ 4.5g/L glucose | Corning | 15-013-CV |
| Penicillin:Streptomycin Solution | Gemini Bio | 400-109 |
| GenClone™ Fetal bovine serum | Genesee Scientific | 25-550 |
| CellGenix® GMP DC | CellGenix | 20901-0500 |
| Recombinant human GM-CSF | PeproTech | 300-23 |
| Recombinant human IL-4 | PeproTech | 200-04 |
| Recombinant human IFNγ | PeproTech | 300-02 |
| LPS | Sigma | L2630 |

### Differentiation of iPSC and ESC to Monocytes

For the control method of iPSC-derived monocyte differentiation, we used the embryoid body (EB) method as described in our earlier work (Taylor et al., 2018). Briefly, iPSC colonies were grown to approximately 5mm in mTeSR1 on Matrigel-coated 6-well plates. Colonies were lifted using a cell lifter and cultured in mTeSR1 supplemented with 10μM ROCK inhibitor (Y-27632) in ultra-low attachement 6-well plates for 4 days to form EBs. After 4 days, EBs were plated on tissue-culture treated 6-well plates in monocyte factory medium (X-VIVO15 medium, supplemented with GlutaMAX^™^, 2-mercaptoethanol, 100ng/mL recombinant human M-CSF, and 25ng/mL recombinant human IL-3). Every 5 days, 2/3 of the culture medium was very carefully aspirated and replaced with fresh monocyte factory medium without disturbing the attaching EBs. Monocyte production generally started after 3-4 weeks.

ESC or iPSC were differentiated to hematopoietic progenitors following the manufacturer’s instructions for the STEMdiff Hematopoietic Kit (STEMCELL Technologies). Briefly, ESC or iPSC were grown to ~75% confluency then dissociated into aggregates using ReLeSR (STEMCELL Technologies) and suspended in mTeSR Plus. ESC or iPSC aggregates were gently passed through an inverted sterile 70μm nylon cell strainer. The cell strainer was reversed and ≥70μm aggregates were gently washed off in 1mL of mTeSR Plus and the clusters counted. Aggregates were plated (~60 per well) in a Matrigel coated 12-well plate in 1mL of mTeSR Plus medium. The following day, it was verified that there were 16-40 colonies attached in each well. Over the next 12 days, the medium was changed according to the STEMdiff Hematopoietic kit instructions and on day 12, CD34^+^ cells were harvested from the plate leaving behind a hemogenic endothelium. The CD34^+^ cells were cryopreserved at 10e6 cells/mL in Cryostor CS10 (STEMCELL Technologies) according to the manufacturer’s protocol and the remaining hemogenic endothelium was cultured in 2mL of monocyte factory medium. The hemogenic endothelium began producing monocytes over the next 8-9 days and continued producing monocytes for 30+ weeks (Figure 1C). Monocytes were harvested with each medium change (every 3-7 days depending on how quickly the medium acidified) by gently rinsing monocytes off each well with the spent medium before passing through a 70μM nylon filter and centrifuging at 300xg for 5 minutes at room temperature (RT). The medium in each harvested well was immediately replaced with 2mL of fresh monocyte factory medium. Cell count, viability, and diameter was measured on a Cellometer Auto 2000 Cell Viability Counter (Nexcelom Bioscience, Lawrence, MA, USA). iPSC and ESC-derived monocytes were used immediately.

### Isolation of Primary Monocytes

Peripheral blood mononuclear cells (PBMCs) were isolated from heparinized whole blood or leukopaks (LifeSouth Community Blood Center, Gainesville, FL) under approval by the Institutional Review Board at the University of Florida. The heparinized whole blood or leukopaks were diluted 1:1 with PBS containing 2mM EDTA. To isolate PBMCs, 25mL of the dilution was overlaid on 15mL of Ficoll-Paque (GE Healthcare, Chicago, IL, USA) and centrifuged at 1200xg for 20 minutes at RT in a swinging-bucket rotor without brake. The buffy coat, containing the PBMCs, was carefully transferred to a new 50mL tube and diluted to 50mL with PBS containing 2mM EDTA and centrifuged at 400xg for 10 minutes at RT. To remove contaminating red blood cells, the pellet was suspended in 5 mL of ACK lysis buffer (ThermoFisher), incubated at RT for 5 minutes, and centrifuged at 300xg for 5 minutes at RT. To remove platelets, the cells were washed three times by aspirating the medium, suspending in 50mL of PBS containing 2mM EDTA, and centrifuging at 400xg for 10 minutes a RT. CD14 MicroBeads (MACS Miltenyi Biotec, Auburn, CA, USA) were used to isolate monocytes from PBMCs according to the manufacturer’s protocol. After isolation the CD14^+^ cells were counted and cell diameter was measured on a Cellometer Auto 2000 Cell Viability Counter. Monocytes were immediately used or cryopreserved at 10e6 cells/mL in CryoStor CS10 according to the manufacturer’s protocol.

### Differentiation of Primary, iPSC, or ESC-derived Monocytes to Macrophages

Monocytes were differentiated to MDM as previously described (Warren et al., 2010). Briefly, freshly isolated primary monocytes, cryopreserved primary monocytes, or freshly harvested iPSC or ESC-monocytes were plated at 4e6 cells per well in a 6-well ultra-low attachment plate in 4mL of MDM medium (DMEM with 4.5 g/L glucose (Corning), 10% heat-inactivated fetal bovine serum (Genesee Scientific, El Cajon, CA, USA), GlutaMAX^™^, and 100 IU penicillin–streptomycin (Gemini Bio, Sacramento, CA, USA)) supplemented with 10ng/mL of recombinant human M-CSF (rhM-CSF) on day 0. On day 2, 2mL of MDM medium supplemented with 10ng/mL of rhM-CSF was added to each well. Between days 7 and 10, monocyte-derived macrophages (MDM) were gently dissociated from the plate using Cell Dissociation Buffer, enzyme-free, PBS (ThermoFisher), pelleted at 300xg for 5 minutes at RT, suspended in fresh MDM medium, and plated for experiments.

### Differentiation of Primary, iPSC or ESC-derived Monocytes to Dendritic Cells

Monocyte differentiation to moDC can be accomplished with the addition of IL-4 and GM-CSF (Taylor et al., 2018). On day 0, freshly isolated primary monocytes, cryopreserved primary monocytes, or freshly harvested iPSC or ESC-monocytes were plated at 4e6 cells per well in a 6-well ultra-low attachment plate (Corning) in 4mL of GMP DC medium (CellGenix, Portsmouth, NH, USA) supplemented with GlutaMAX^™^, 50ng/mL recombinant human IL-4 (rhIL-4 [PeproTech]), and 50ng/mL recombinant human GM-CSF (rhGM-CSF [PeproTech]). On day 2, 2mL of GMP DC medium supplemented with GlutaMAX^™^, 50ng/mL rhIL-4, and 50ng/mL rhGM-CSF was added to each well. On days 4 and 6, 2mL of medium was carefully removed from each well and replaced with 2mL of fresh GMP DC medium supplemented with GlutaMAX^™^, rhIL-4 (50ng/mL), and rhGM-CSF (50ng/mL). On day 7, immature monocyte-derived DCs (moDC) were gently washed off the plate, centrifuged at 300xg for 5 minutes at RT, suspended in fresh GMP DC medium supplemented with GlutaMAX^™^, and plated for experiments. To mature moDC, 200ng/mL of LPS (Sigma, St. Louis, MO, USA) and 50ng/mL recombinant human IFNγ (rhIFNγ [PeproTech]) were added on day 6.

### Morphology of Primary, iPSC, and ESC-derived Monocytes, MDM, and moDC

For primary, iPSC, or ESC-derived monocytes, freshly isolated cells were immediately fixed/permeabilized for staining using the Fixation/Permeabilization Solution Kit at 4°C for 20 minutes. MDM were plated at 5e5 cells per dish on a 35mm glass bottom microwell dish (MatTek, Ashland, MA, USA) in 4mL of MDM medium. Cells were allowed to attach and then fixed/permeabilized using the Fixation/Permeabilization Solution Kit at 4°C for 20 minutes. moDC were washed off the plate following differentiation then fixed and permeabilized like monocytes. All fixed/permeabilized cells were washed three times in room temperature Perm/Wash buffer and then stained in 2mL of Perm/Wash buffer with 4 drops of the NucBlue Fixed Cell ReadyProbes reagent (ThermoFisher) for 15 minutes at room temperature. Monocytes and MDM were also stained with 4 drops of the ActinGreen 488 ReadyProbes reagent (ThermoFisher). Cells were gently rinsed twice with Perm/Wash buffer and 1.5mL of Live Cell Imaging Solution (ThermoFisher) was added. Primary and iPSC-derived cells were imaged with an EVOS FL Imaging System (ThermoFisher) using the EVOS DAPI (Excitation: 470/22nm, Emission: 510/42nm) and GFP (Excitation: 357/44nm, Emission: 447/60nm) light cubes (ThermoFisher). ESC-derived cells were imaged on a Leica Fluorescent DMI 6000B wide field microscope (Leica, Wetzlar, Germany).

### Surface Marker Expression of Primary, iPSC, and ESC-derived Monocytes, MDM, and moDC

Cultured cells were pelleted at 300xg for 5 minutes at RT and suspended in flow staining buffer (PBS containing 5% FBS and 2mM EDTA). Fc receptors were blocked for 10 minutes at 4°C with FcR blocking reagent (MACS Miltenyi Biotec). Cells harvested during the conversion period from CD34^+^ hematopoietic precursors to CD14^+^ monocytes (Days 12-21 of the differentiation protocol) were stained with αCD14-PE (BioLegend, San Diego, CA, USA) and αCD34-APC (BioLegend). Monocytes were stained with αCD14-PE, αCD16-FITC (ThermoFisher), and αCD64-APC (ThermoFisher). MDM were stained with αCD11b-FITC (BioLegend) and αCD68-PE (ThermoFisher). Primary and iPSC-derived moDC were stained with αCD11c-APC (BioLegend), αHLA-DR, DP, DQ-APC (BioLegend), and αCD209-APC (ThermoFisher). ESC-derived moDC were stained with αCD11c-FITC (BioLegend), αHLA-DR, DP, DQ-PE/Cy7 (BioLegend), and αCD209-APC (ThermoFisher). All antibody information is in Table 2. Surface markers were stained for 30 minutes at 4°C. Surface-stained cells were washed 3 times in flow staining buffer and analyzed immediately on an Accuri C6 flow cytometer. Surface stained primary and iPSC-derived cells were fixed/permeabilized and stained using the Fixation/Permeabilization Solution Kit according to the manufacturer’s instructions. Surface-stained ESC-derived cells were fixed/permeabilized and stained using the eBioscience FOXP3/Transcription Factor Staining Buffer Set (ThermoFisher) according to the manufacturer’s instructions. Primary and iPSC-derived cells were analyzed on an Accuri C6. ESC-derived cells were analyzed on a LSRII (BD Biosciences). All data were analyzed in FlowJo Version 10.

### MDM Phagocytosis, Endocytosis, and Peptide Processing Assays

Primary or iPSC-derived MDM were plated at 2e5 cells per well in a 24-well ultra-low attachment plate in 0.5mL of MDM medium. Cells were allowed to rest/attach for 12 hours. After 12 hours, 250μL of medium was removed and replaced with 250μL of fresh medium supplemented with 100μg/mL pHrodo Red *S. aureus* bioparticles (ThermoFisher), to track phagocytosis, pHrodo Red dextran (ThermoFisher), to track endocytosis, or DQ-Ovalbumin (ThermoFisher) to track peptide processing for loading onto MHC and incubated at 37°C on a time course. At end point, MDM were gently washed twice with 1mL of 37°C PBS, 0.5mL of 4°C Cell Dissociation Buffer was added to each well, and the plate was incubated on ice for 5 minutes. MDM were gently washed off the plate and pelleted at 500xg for 5 minutes in a 4°C centrifuge. MDM were suspended in 100μL of 4°C annexin V binding buffer (BioLegend) and the two pHrodo loaded sets were stained with 2μL of FITC Annexin V (BioLegend), while the DQ-Ovalbumin loaded set was stained with APC Annexin V (BioLegend) for 5 minutes at RT. After 5 minutes, 100μL of annexin V binding buffer was added and cells were immediately analyzed on an Accuri C6 (BD Biosciences). Phagocytosis of pHrodo Red *S. aureus* bioparticles, endocytosis of pHrodo Red dextran, and proteolysis of DQ-Ovalbumin were analyzed immediately on an Accuri C6. Data were analyzed in FlowJo version 10 software.

### moDC Phagocytosis, Endocytosis, and Peptide Processing Assays

Primary or iPSC-derived moDC were plated at 2e5 cells per well in a 96-well round bottom plate in 0.1mL of GMP DC medium. After plating, 5μL of 1000μg/mL pHrodo Red *S. aureus* bioparticles, pHrodo Red dextran, or DQ-Ovalbumin was added to each well on a time course and incubated at 37°C until the experimental end point. At end point, 100μL of 4°C annexin V binding buffer containing 2μL of FITC Annexin V was added to the two pHrodo loaded sets and 100μL of 4°C annexin V binding buffer containing 2μL of APC Annexin V was added to the DQ-Ovalbumin loaded set and cells were stained on ice for 5 minutes. Phagocytosis of pHrodo Red *S. aureus* bioparticles, endocytosis of pHrodo Red dextran, and proteolysis of DQ-Ovalbumin were analyzed immediately on an Accuri C6. Data were analyzed in FlowJo version 10 software.

### Macrophage Activation

Primary or iPSC-derived MDM were plated at 2e5 cells per well in a 24 well plate in 500μL of MDM medium. Cells were left untreated or activated with 100ng/mL of LPS, 10ng/mL of rhIFNγ, or both LPS and r IFNγ hIFNγ. After 48 hours of activation, supernatants were harvested and assayed for human IL-10, TNF, and CXCL10 via enzyme linked immunosorbent assay (ELISA) (ThermoFisher). ELISA data was acquired on a SpectraMax M5 Multi-Mode Microplate Reader (Molecular Devices, San Jose, CA, USA).

### Dendritic Cell/CD8^+^ T Cell Co-culture

Primary or iPSC-derived moDC were plated at 0.1e6 cells per well in a 96-well round bottom plate in 0.1mL of GMP DC medium and allowed to rest for 48 hours. After 48 hours, moDC were gently washed twice with T cell expansion medium (RPMI supplemented with HyClone FBS [GE Healthcare, Chicago, IL, USA], 1000U/mL Penicillin/streptomycin, Sodium Pyruvate [Corning], HEPES [Corning], nonessential amino acids [Corning], GlutaMAX^™^, and β-mercaptoethanol) and loaded with MART-1 peptide (ELAGIGILTV; thinkpeptides, Littlemore, Oxford, UK) at 50μM or left unloaded and incubated for 2 hours. After 2 hours, Carboxyfluorescein succinimidyl ester (CFSE)-stained MART1-CD8 T cell avatars were added at the T cell:APC ratio of 3:1 (0.3e6 cells per well) in 0.2mL of complete T cell medium. Cells were co-cultured for 5 days then CFSE staining was analyzed on an Accuri C6 flow cytometer. Data were analyzed in FlowJo version 10 software.

