## Supplementary figures and images for "High-Yield Monocyte, Macrophage, and Dendritic Cell Differentiation from Induced Pluripotent Stem Cells"

### Supplemental Figure 1

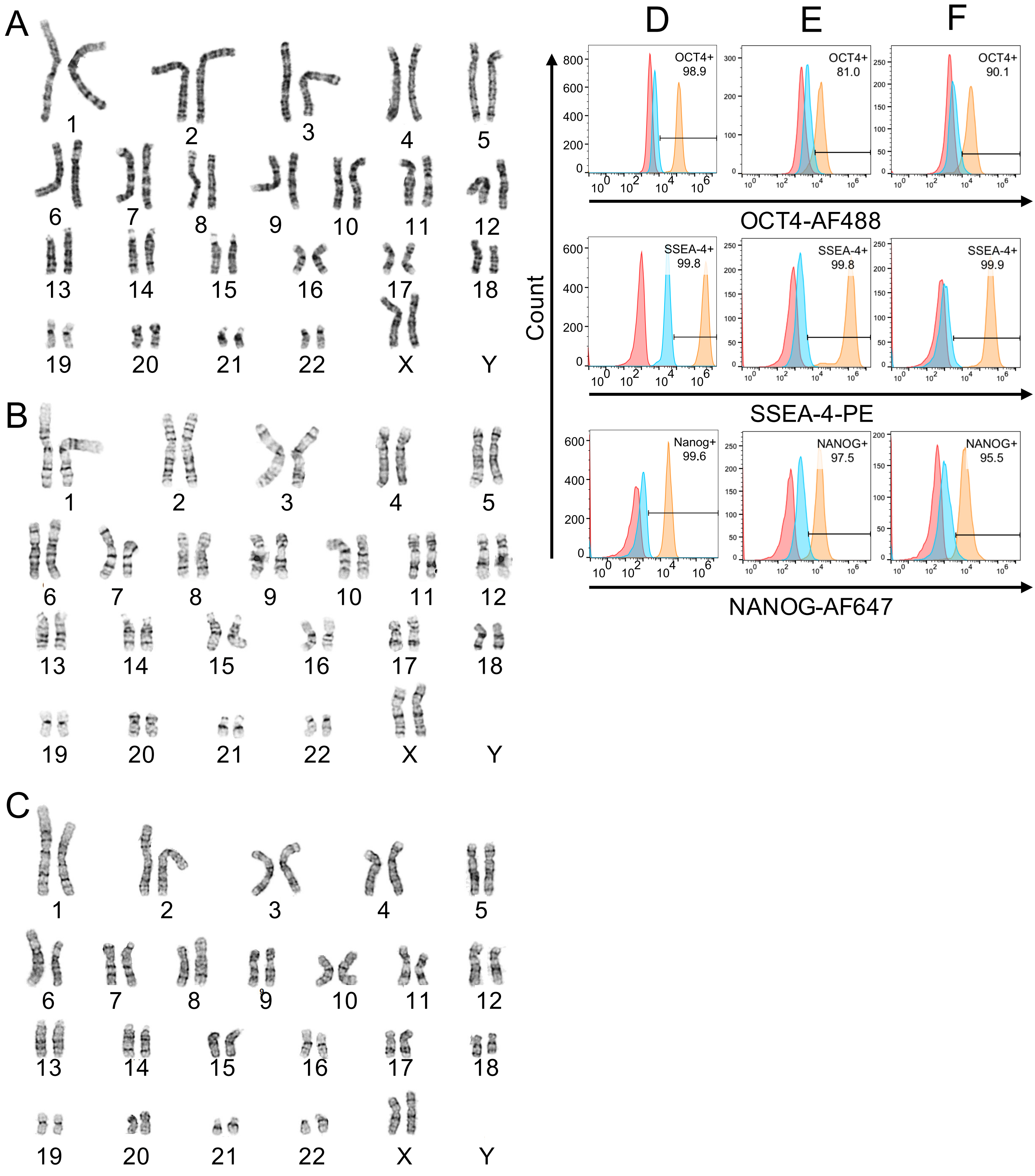

### Supplemental Figure 2

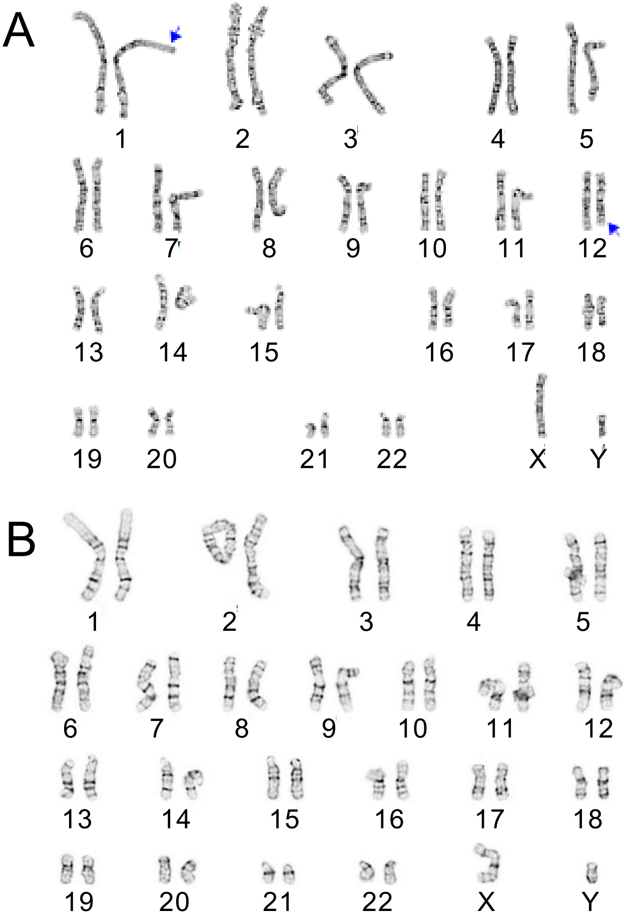
